# Long term nasopharyngeal colonization by *Staphylococcus aureus* determinants of adaptation

**DOI:** 10.1101/2021.09.19.460983

**Authors:** Breno A. B. Salgado, Elaine M. Waters, Josephine C. Moran, Aras Kadioglu, Malcolm J. Horsburgh

## Abstract

*Staphylococcus aureus* nasal colonization is a risk factor for infection. A large proportion of the population are identified as potential *S. aureus* carriers yet we only partially understand the repertoire of genetic factors that promote long-term nasal colonization. Here we present a novel murine model of nasopharyngeal colonization that requires a low *S. aureus* inoculum and is amenable to experimental evolution approaches. We used this model to experimentally evolve *S. aureus* using successive passages in the nasopharynx to identify those genetic loci under selection. After 3 cycles of colonization, mutations were identified in mannitol, sorbitol, arginine, nitrite and lactate metabolism genes promoting key pathways in nasal colonization. Stress responses were identified as being under selective pressure, with mutations in DNA repair genes including *dnaJ* and *recF* and key stress response genes *clpL, rpoB* and *ahpF*. Peptidoglycan synthesis pathway genes also revealed mutations indicating potential selection for alteration of the cell surface. The infection model used here is versatile to assist decolonization and persistence studies.

## Introduction

*S. aureus* is an opportunistic pathogen commonly found in both the anterior and inner nasal cavity. While *S. aureus* carriage is typically asymptomatic, it is a risk factor for infection. Since the eradication of carriage prevents both infection and transmission of *S. aureus* [1-3], knowledge of the determinants that enable *S. aureus* to colonize the human nasopharynx are critical in developing strategies to control the spread of *S. aureus*.

The major components contributing to *S. aureus* nasal colonization depends on the composition of the niche [4]. The anterior nares are lined with keratinized epithelium identical to skin, with loricrin, K1 and K10 cytokeratin molecules present in high numbers. *S. aureus* cell wall-anchored (CWA) proteins, such as clumping factor B (ClfB) and iron-regulated surface determinant A (IsdA) are major adhesins contributing to successful colonization of the anterior nasal cavity [5]. Other CWA proteins, such as SdrC, SdrD and SasG, are also associated with *S. aureus* nasal colonization [6]. The epithelium of the inner nasal sites where *S. aureus* is commonly found is characterized by a ciliated and pseudostratified morphology. The cells in this niche express SREC-I, a member of the F-type scavenger receptor family, a key component in *S. aureus* colonization recognized by *S. aureus* glycopolymer wall teichoic acid (WTA) [7,8]. While the interaction between *S. aureus* adhesins and its ligands in the anterior nares are implicated in persistent colonization, the WTA/SREC-I interface is proposed to be important in the early stages of nasal colonization [4].

During long-term colonization, *S. aureus* is exposed to diverse selective pressures that include competing microbiota, the host immune system and nutrient limitation [9.10]. *S. aureus* must therefore be versatile in order to persist, and is known for its high evolutionary rate, resulting in large genomic and phenotypic diversity [11,12]. Genetic variation from mutations in the form of single nucleotide polymorphisms (SNPs) or short insertion/deletions (INDELs) can be tracked by whole-genome sequencing; a method previously used to identify contributory genes involved in *S. aureus* adaptation to specific selective pressures over an evolutionary time-frame [13,14]. For example, comparative genomics was used to detect *S. aureus* genes and alleles with a role in bacteraemia [15] and infections during cystic fibrosis [14]. However, detailed temporal mechanisms of nasal colonization remain unclear with several determinants of *S. aureus* carriage yet to be characterized, highlighting the need for robust models that mirror more reliably *in vivo* conditions [16].

Experimental evolution of microbes selects for adaptation to applied environmental pressure and the use of whole-genome sequencing simplifies the identification of genomic adaptations acquired by this directed evolution. Several studies described *S. aureus* evolution *in vitro* by serially passaging isolates over days or even weeks under selective pressure from antimicrobials to reveal adaptation and survival mechanisms [17-19], however there are no reports of equivalent studies using *in vivo* models.

Here we developed an *in vivo* long-term asymptomatic nasopharyngeal colonization model for the experimental evolution of *S. aureus* to investigate temporal colonization of the mouse nasopharynx. The isolates generated using this approach were sequenced to identify genetic changes that might reveal key components, pathways and mechanisms that enable *S. aureus* to colonize and persist in the nasal cavity.

## Results

### Establishing a novel long-term *S. aureus* nasopharyngeal colonization model for experimental evolution

A long-term *S. aureus* nasopharyngeal colonization model was established by inoculating groups of mice intranasally with ∼ 5 × 10^4^ CFU *S. aureus* USA300 LAC JE2 strain. The mice were maintained for a period of 14 days post-infection. At chosen time points (1, 3, 5, 7 and 14 days post-infection), the nasopharyngeal CFU load in each mouse was determined (Fig 1).

**Figure 1.**
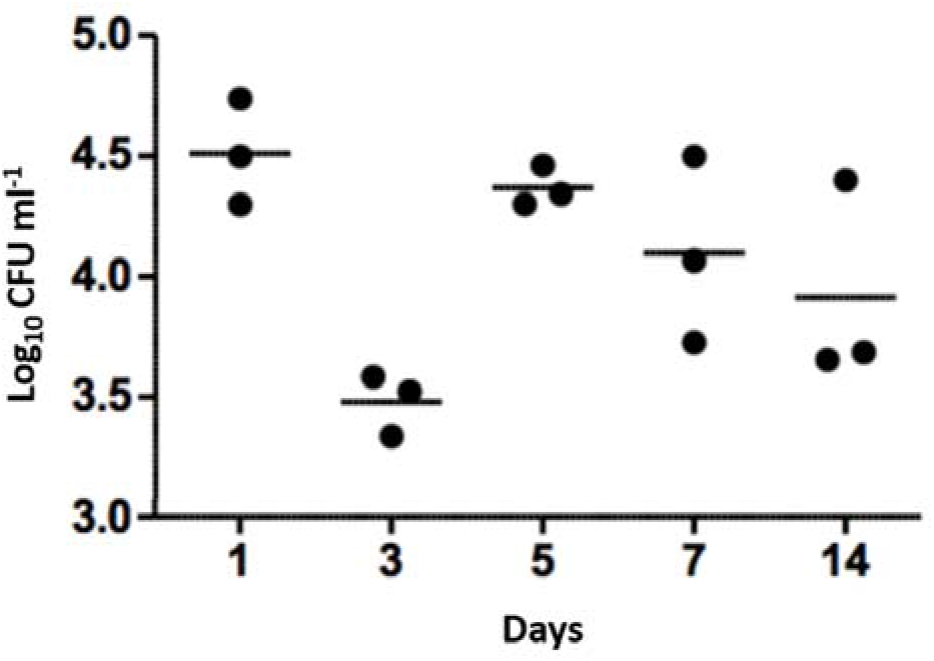
Nasopharyngeal colonization of mice by *S. aureus* USA300 LAC JE2. Intranasal inoculation with ∼5 × 10^4^ CFU. At each time point post infection, the colonizing bacteria were recovered and cultured to identify *S. aureus*. Bars represent mean values of the mice in each time group.

Across the 14 days of nasopharyngeal colonization, numbers of viable *S. aureus* CFU decreased by ∼1 log by day 3 compared to day 1, followed by an increase in CFU from day 5 onwards, with the population remaining constant thereafter until day 14 when the experiment was ended (Fig 1). The model was therefore determined to be sufficiently robust to design an experimental evolution using serial passages.

### Repeated nasopharyngeal colonization selects mutations in *S. aureus*

Based upon the established colonization model, 3 serial passages of *S. aureus* were performed in total with each passage lasting for 7 days (Fig 2). Isolated bacteria from the first passage were used as inoculum for the second passage. The process was repeated using the bacteria from the second passage as inoculum for the third passage. The three mice from each nasopharyngeal passage survived over the incubation period up until they were sacrificed to recover the colonizing *S. aureus*. After completion of the serial passages, *spa* typing of the bacteria was performed by DNA sequencing of the amplified gene to confirm nasopharyngeal isolates matched the input strain.

**Figure 2.**
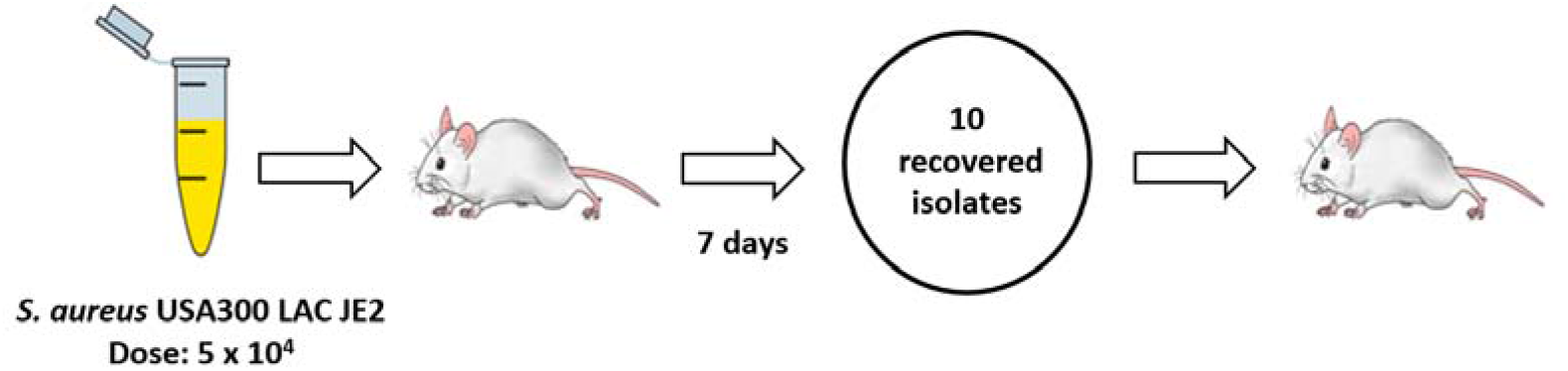
Serial passage of *S. aureus* in a murine nasopharyngeal colonization model. Schematic demonstration of sequential colonization of mouse nasopharynx by *S. aureus* USA300 LAC JE2. Three passages were performed with each lasting 7 d. An inoculum of ∼5×10^4^ CFU was used for the first passage. For the second and third passages 10 isolates randomly collected from the previous passage were pooled and adjusted to ∼5×10^4^ CFU to serve as inoculum. Three mice were used per round of colonization.

Whole genome sequencing of 10 pooled isolates from each mouse in the three passages was used to determine whether repeated *S. aureus* colonization of the nasopharynx selected for genetic changes. The isolate inputs for pooled for DNA sequencing are shown in Table 1.

**Table 1.** List of strains used and generated in this study.

| Origin | Description | Reference |
| --- | --- | --- |
| **Control** | *S. aureus* USA300 LAC JE2 (1 isolate) | Fey et al., 2013 |
| **First Passage** | Mouse A1 (10 isolates) | This study |
|  | Mouse B1 (10 isolates) | This study |
|  | Mouse C1 (10 isolates) | This study |
| **Second Passage** | Mouse A2 (10 isolates) | This study |
|  | Mouse B2 (10 isolates) | This study |
|  | Mouse C2 (10 isolates) | This study |
| **Third Passage** | Mouse A3 (10 isolates) | This study |
|  | Mouse B3 (10 isolates) | This study |
|  | Mouse C3 (10 isolates) | This study |

After the first passage, there was determined to be limited intragenic and intergenic sequence variation (Fig 3) based upon a threshold imposed of variants being represented at >20% i.e in at least two isolates. Seven SNPs in total were identified from the 3 mice, with only one SNP being observed in more than one pool of isolates (S1 and S2 Tables). Identified SNPs showed low frequency of around 20% of total reads, indicating that at least two of the ten isolates forming the pooled genomic DNA carried the identified mutation. One sequenced isolate DNA pool (mouse B1) did not bear any SNPs. Overall, the low abundance and frequency of SNPs suggests there was not a pronounced or uniform selection on *S. aureus* during the first seven days of nasopharyngeal colonization as evidenced by the lack of a dominant variant causing population change.

**Figure 3.**
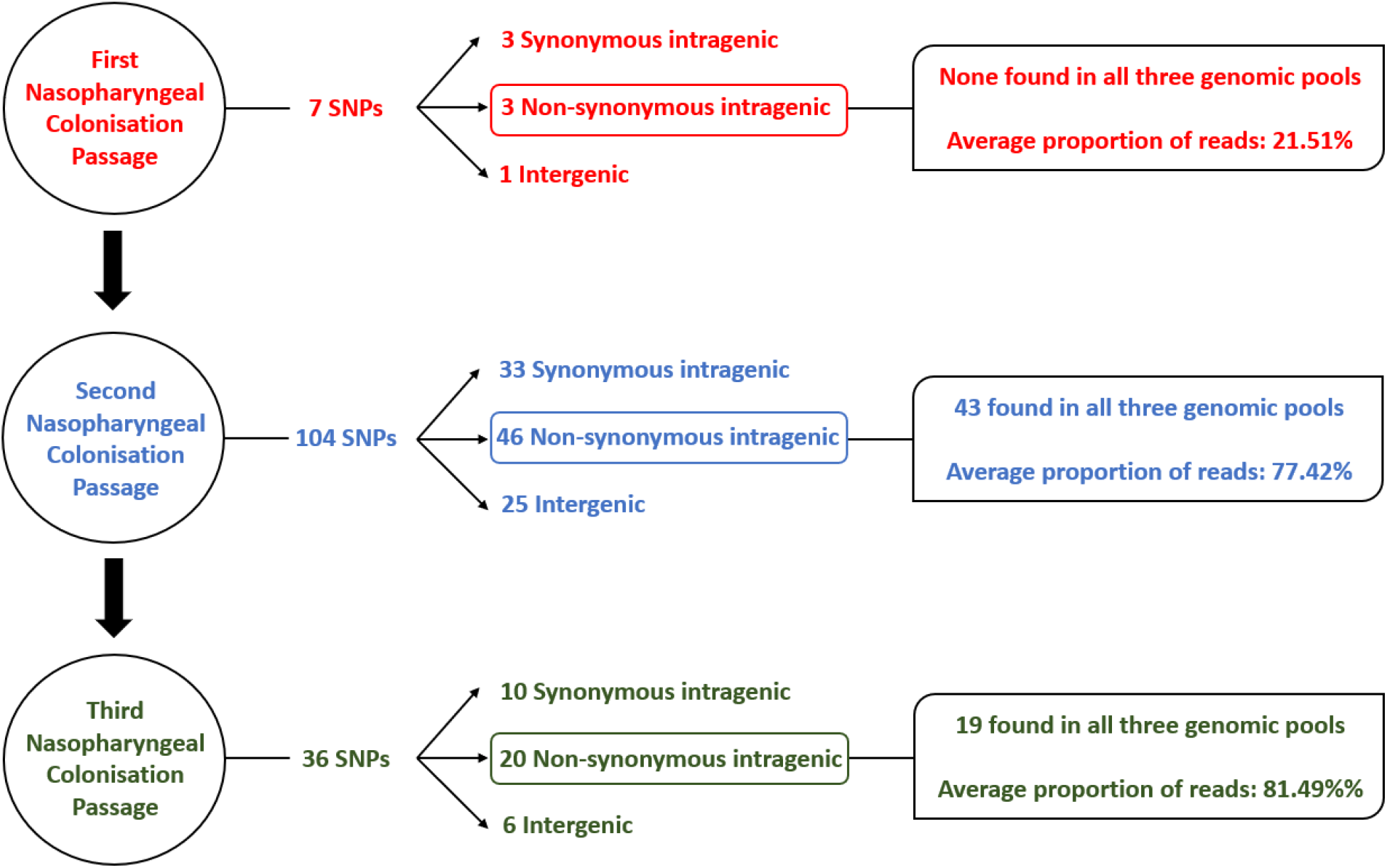
Flow chart detailing sequence variants identified in *S. aureus* isolates after each nasopharyngeal passage.

DNA sequencing of isolates after the second nasopharyngeal passage revealed a greater number of variants in the *S. aureus* clones, with 104 SNPs found across the three genomic DNA pools (Fig 3) based on the same >20% threshold imposed in the first round of sequence analysis. From these variants, 43 out of 46 non-synonymous intragenic SNPs and 29 out of 30 intergenic SNPs were observed in all three genomic pools (S3 and S4 Tables). A further difference between the profiles of sequence variants from the first and second passages is the higher proportion of reads for each variant in the second round. The DNA sequence pools from the second passage (B2 and C2) (Fig 2; S3 Table) revealed 37 intragenic SNPs that were present in at least 90% of the reads, (i.e. in 9/10 of the recovered *S. aureus* isolates). The pool of isolates from A2 had a lower proportion of reads for sequence variants, where most of the SNPs (n=36; 78.2%) were detected in a lower proportion (60%) of the reads, though still substantially higher than the variant frequencies of the first passage. The increased number of sequence variants identified after the second passage was suggestive of selection *S. aureus* during nasopharynx colonization leading to a third passage.

A reduced range of sequence variants was identified across the isolate pools from the three mice after the third 7-day colonization of the nasopharynx passage. A total of 36 SNPs were detected from the genome DNA pools (Fig 3). Above the threshold, sequence variants were present at ∼90% of reads shared across the three sequence pools, with the exception of those detected in the hypothetical protein SAUSA300_RS09205. Among the 20 non-synonymous SNPs detected in the sequence pools of the third passage, 16 persisted from the first and/or second passage. In addition, three out of six intergenic SNPs also persisted across nasopharyngeal passages (Table 2).

**Table 2.** SNPs of nasopharyngeal colonisation isolates that persisted across repeated passages. *S. aureus* genome sequence variants (SNPs) in coding and non-coding regions that appeared in the first and/or second nasopharyngeal passage and was further detected after a third repeated passage in an experimental colonisation model.

| Genome change | Genome symbol | Gene function | Codon change |
| --- | --- | --- | --- |
| 5010 (C/G) | *recF* | DNA replication and repair | 358 aa (P/A) |
| 61025 (G/A) | SAUSA300_RS00265 | Hypothetical protein | 55 aa (L/F) |
| 240358 (T/A) | SAUSA300_RS01070 | Peptide ABC transporter protein | 358 aa (N/K) |
| 270510 (T/C) | SAUSA300_RS01200 | FadB; Hydroxyacyl Coenzyme A | 601 aa (T/A) |
| 292738 (A/C) | *gutB* | Sorbitol dehydrogenase | 341 aa (T/P) |
| 1533312 (A/T)* |  |  |  |
| 376057 (C/T) | SAUSA300_RS01715 | Flavin oxidoreductase | 258 aa (D/N) |
| 376231 (T/G) | SAUSA300_RS01715 | Flavin oxidoreductase | 200 aa (S/R) |
| 460966 (G/A) | SAUSA300_RS02185 | Staphylococcal superantigen-like 11 | 110aa (A/T) |
| 1286986 (A/G) | *ftsK* | DNA translocase | 180 aa (K/R) |
| 1857089 (C/T) | SAUSA300_RS09205 | Hypothetical protein | 179 aa (V/I) |
| 1857109 (A/G) | SAUSA300_RS09205 | Hypothetical protein | 172 aa (I/T) |
| 1857182 (G/A) | SAUSA300_RS09205 | Hypothetical protein | 148 aa (P/S) |
| 1857188 (C/T) | SAUSA300_RS09205 | Hypothetical protein | 146 aa (E/K) |
| 1857202 (G/A) | SAUSA300_RS09205 | Hypothetical protein | 141 aa (A/V) |
| 1857215 (T/C) | SAUSA300_RS09205 | Hypothetical protein | 137 aa (T/A) |
| 1857232 (T/C) | SAUSA300_RS09205 | Hypothetical protein | 131 aa (K/R) |
| 1957938 (C/T)* |  |  |  |
| 2487639 (A/G)* |  |  |  |

\*Intergenic SNP

### Nasopharyngeal colonization-selected SNPs in *S. aureus*

The functional category of genes containing sequence variants after each of the three 7-day nasopharyngeal colonizations were identified. The variants included non-synonymous intragenic SNPs, and intergenic sequence variants determined to have the potential to alter transcription of downstream genes. The majority of the functional categories belong to genes encoding proteins involved in cell metabolism (amino acid and co-factors biosynthesis; energy and fatty acid metabolism) or DNA metabolism (replication, recombination and repair) (Fig 4).

**Figure 4.**
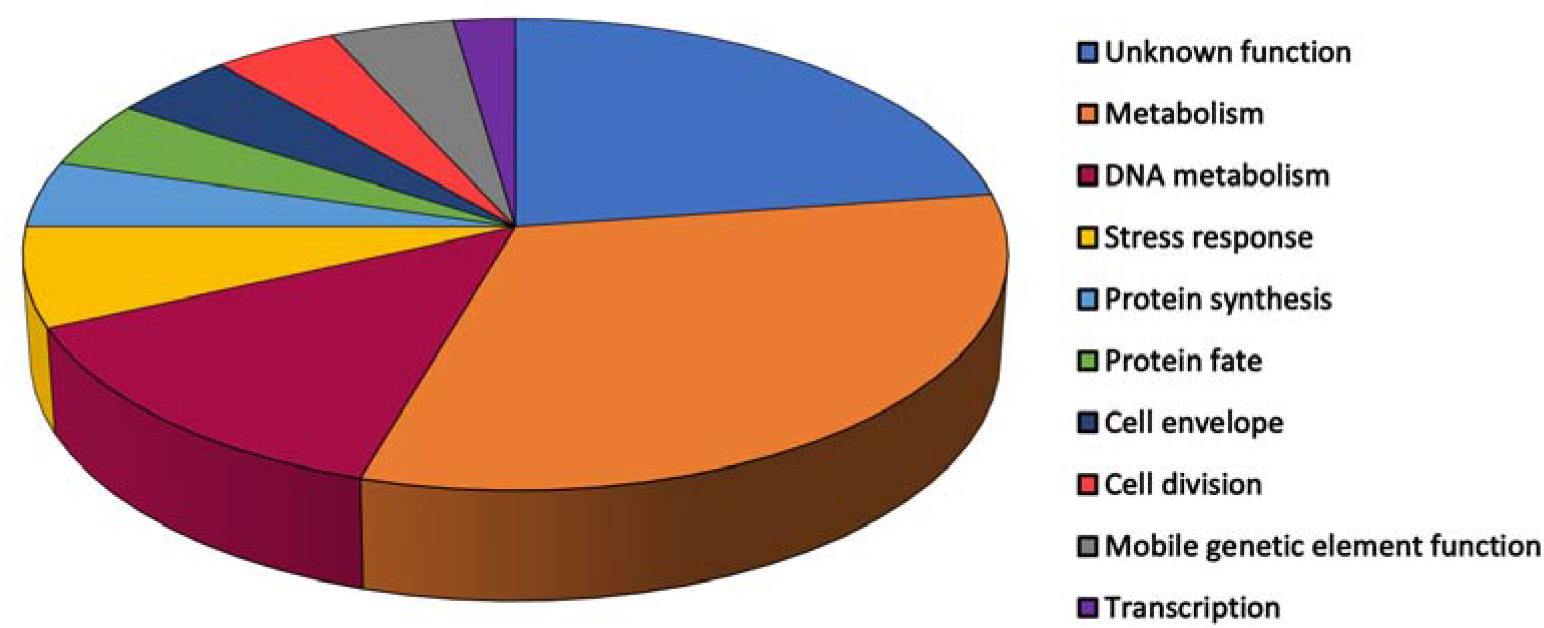
Relative proportions of functional groups ascribed to *S. aureus* genes with identified sequence variants across three serial nasopharyngeal passages. Variants were included for non-synonymous intragenic SNPs (n=44).

Proteins in pathways of DNA replication, recombination and repair with intergenic SNPs included: topoisomerases (GyrA, GyrB, GyrC); translocases (FtsK); nuclease (SbcC), and the DNA replication and repair protein RecF. The relative proportion of genes associated with these DNA pathways harboring SNPs could reflect selection for genome adaptation. Increased mutability by *S. aureus* was reported as an adaptation mechanism [20], whereby weakened DNA replication control and repair promotes allelic variability, benefiting *S. aureus* in its niche.

Variants associated with transcriptional regulation were observed for genes encoding proteins involved in transcription (RpoB), RNA modification (Sun, SAUSA300_RS04890 and SAUSA300_RS09885) and RNA decay (SAUSA300_RS06375) (S3 and S5 Tables). These genes play a role in cellular pathways that modulate ribosomal stability and the synthesis of proteins [21] as part of mechanisms the bacteria use to changes in the microenvironment [22].

Sequence variants were identified in genes encoding metabolism proteins including mannitol (MtlA) and sorbitol fermentation (GutB), fatty acid metabolism (FadB; SAUSA300_RS01200 and SAUSA300_RS07925) and peptide metabolism (AmpA aminopeptidase). Further sequence variants were identified in genes encoding proteins associated with the biosynthesis of co-factors biotin (BioA) and thiamine (SAUSA300_RS11285), haem (SAUSA300_RS08410), and amino acids valine (ValS) and histidine (HisG) following *S. aureus* nasopharyngeal colonization. Genes encoding proteins within arginine metabolism revealed variants: ArcA and ArgF. Both enzymes catalyze formation of citrulline, the former as part of the arginine deiminase pathway that serves as energy source for *S. aureus* in anaerobic conditions, and the latter within arginine synthesis from glutamate [23,24]. Pathways for responses to a variety of stresses revealed sequence variants within genes encoding ClpL (general stress tolerance), AhpF (oxidative stress) and DnaJ (hyperosmotic and heat shock stresses) together with variants observed in genes associated with anaerobic metabolism (NarK) and lactate (LctP2).

Those sequence variants that were identified within the pathways described during nasopharyngeal colonization revealed a smaller subset that persisted across the successive rounds of colonization (Table 2). One gene was identified as the target for multiple mutations across its length with varied individual frequencies of the variants from the three mice in the final passage. The gene (SAUSA300_RS09205) encodes a 399 amino acid hypothetical membrane protein described within the TIGRFAM: transport proteins and PFAM: metallopeptidase family M24. The collective sequence variants identified reveal this gene as showing selection most clearly across the sequential passages. Conversely, from the experimental evolution sequence data, there were sequence variants not maintained temporally, indicating potential refinement of selection during the repeated nasopharyngeal colonization passages of *S. aureus*. The vast majority of selected variants that persisted across repeated nasopharyngeal passages of *S. aureus* are genes involved in metabolism and DNA replication (Table 2).

### Resident *S. aureus* prevent colonization of the murine nasopharynx

Experiments to perform the third nasopharyngeal passage repeatedly returned *S. aureus* with a different *spa* type from the USA300 LAC species inoculum. indicating that sets of mice were colonized by *S. aureus* upon arrival. Recent studies reported there is frequent colonization of laboratory mice by *S. aureus* [25] in contrast to earlier studies that *S. aureus* are not natural colonizers of mice [26,27]. Others have determined that colonization by *S. aureus* prevents the invasion of distinct *S. aureus* strains [28]. To gain knowledge of the resident strain in the purchased mice and enable a simple comparative interrogation, we performed whole-genome sequencing and variant analysis on the mouse resident *S. aureus* (named SA_MOU). We sought insights of genome sequence polymorphism that converged with the experimental evolution selection from repeated colonization of the nasopharynx.

Multilocus sequence typing technique (MLST) identified SA_MOU as ST8, the same sequence type of USA300. A recent study demonstrated that the genetic variation within ST8 clones is rather substantial temporally and globally [29] and that ST8 possesses clonal heterogeneity in Europe, where it co-exists with the dominant ST80, and others such as ST22 [30]. The sequenced and assembled SA_MOU genome was aligned with the USA300 reference genome plus representatives of dominant STs in Europe: ST22 and ST80. Alignment showed fewest gaps with the SA_MOU assembled contig against the USA300 reference genome, supported by a pairwise identify of 99.5% across 94.5% coverage as the highest values among the three genome alignments (data not shown). Alignment of SA_MOU against ST22 and ST80 genomes revealed high pairwise identify (97.9% and 99.4%, respectively) but with lower coverage (88.0% and 90.9%, respectively).

Because we identified that SA_MOU was resident in mouse nasopharynx and was the same ST as USA300 LAC JE2, we hypothesized there was concordance between sequence variants that occurred in passaged isolates and the corresponding SA_MOU allele. Matches could provide support for key loci undergoing *in vivo* selection. The genes containing 13 non-synonymous intragenic SNPs from the third experimental nasopharyngeal passage were compared with those of the SA_MOU annotated genome. Sequences for 9 of the 13 genes were identified in the SA_MOU genome (Table 3) with four loci not located in the assembled genome. The cognate protein sequences detected in SA_MOU were compared with the translated USA300 reference genome (NC_007793.1) using BLASTp, which identified two proteins with matched sequence variation: RecF and FtsK.

**Table 3.** Non-synonymous intragenic SNPs comparison between two ST8 *S. aureus* nasopharyngeal strains. SNPs observed in genomic pools (3 × 10 isolates) from colonisation isolates that persisted across three repeated nasopharyngeal passages (USA 300 LAC JE2) and a presumed nasopharynx mouse-adapted strain (SA_MOU; 10 isolates).

| Genome change | Strain | Codon change |
| --- | --- | --- |
| *recF* | *S. aureus* USA 300 LAC JE2<br>*S. aureus* SA_MOU | 358 aa (P/A)<br>358 aa (P/A) |
| SAUSA300_RS00265 | *S. aureus* USA 300 LAC JE2<br>*S. aureus* SA_MOU | 55 aa (L/F)<br>No variation |
| SAUSA300_RS01070 | *S. aureus* USA 300 LAC JE2<br>*S. aureus* SA_MOU | 358 aa (N/K)<br>No variation |
| SAUSA300_RS01200 | *S. aureus* USA 300 LAC JE2<br>*S. aureus* SA_MOU | 601 aa (T/A)<br>No variation |
| *gutB* | *S. aureus* USA 300 LAC JE2<br>*S. aureus* SA_MOU* | 341 aa (T/P)<br>- |
| SAUSA300_RS01715 | *S. aureus* USA 300 LAC JE2<br>*S. aureus* SA_MOU | 258 aa (D/N) and 200 aa (S/R)<br>No variation |
| SAUSA300_RS02185 | *S. aureus* USA 300 LAC JE2<br>*S. aureus* SA_MOU* | 110 aa (A/T)<br>- |
| *ftsK* | *S. aureus* USA 300 LAC JE2<br>*S. aureus* SA_MOU | 180 aa (K/R)<br>180 aa (K/R) |
| *dnaJ* | *S. aureus* USA 300 LAC JE2<br>*S. aureus* SA_MOU | 250aa (A/T)<br>No variation |
| SAUSA300_RS09205 | *S. aureus* USA 300 LAC JE2<br>*S. aureus* SA_MOU | 172 aa (I/T); 148 aa (P/S);<br>146 aa (E/K); 141 aa (A/V);<br>137 aa (T/A) and 131 aa (K/R)<br>No variation |
| SAUSA300_RS15385 | *S. aureus* USA 300 LAC JE2<br>*S. aureus* SA_MOU* | 28 aa (Y/C)<br>- |
| SAUSA300_RS10080 | *S. aureus* USA 300 LAC JE2<br>*S. aureus* SA_MOU* | 48 aa (R/L)<br>- |
| *bioA* | *S. aureus* USA 300 LAC JE2<br>*S. aureus* SA_MOU | 189 aa (R/H)<br>No variation |

\*Locus not identified in the assembled SA\_MOU genome

## Discussion

In this study, a novel *S. aureus* nasopharyngeal colonization model was established in mice to enable experimental evolutionary approaches to reveal contributory genetic factors. The colonization model used *S. aureus* USA300 LAC JE2 and showed relatively stable colonisation of the bacteria in the nasopharynx over a 14 day post-inoculation period. Based on these data, day 7 was selected as a timepoint for the design of sequential colonizations that would allow stable carriage and isolation of *S. aureus*.

The colonization data that was determined here was broadly similar to that of a previously developed cotton rat nasal colonization model [31]. A range of different animal models exist for the study of *S. aureus* nasal carriage where an inoculum of >10^7^ CFU are typically used for prolonged colonization [32-36]. Several models use antibiotic pretreatment to promote stable nasal carriage, showing colonization >7 d being achieved with streptomycin-treated mice [27,32]. In all of these models though, the infecting inoculum is high (>10^7^ CFU) which is distinctly dissimilar to natural human colonization. Furthermore, such high infective doses can disrupt the host barrier, significantly altering host-pathogen interactions, while the use of antibiotics can disrupt the native flora potentially altering the host niche. The novel model described here uses a relatively low inoculum of *S. aureus* (5 × 10^4^ CFU), more in keeping with natural colonisation, and which resulted in long term stable carriage without antibiotic pretreatment.

After a first round of colonizations lasting 7 d in our study, DNA sequencing of 10 pooled isolates revealed a low level of genetic variance with only 3 SNPs identified above a set threshold of 20% of sequencing reads. Two subsequent passages of *S. aureus* in the nasopharynx resulted in considerably greater genetic variation. The basis for the expansion of genetic variants by selection in these passages was not identified but could arise from the genetic variants present below the 20% threshold set for calling SNPs after the first round of colonizations or the SNPs present above the threshold increasing mutation rate. Across the three rounds of colonization, there was evidence of genetic variants predominantly being within the DNA repair, recombination and replication category that might signify pathways through which selection has occurred sequentially. The nasopharyngeal colonization model demonstrates a platform to design experimental evolution over a short evolutionary timescale and could be modified to remove the population bottleneck that was imposed with each passage by choosing a limited number of isolates as the input inoculum.

A diverse set of loci and gene functions showed sequence variation as a result of the experimental evolution and reveal determinants that could contribute to the establishment and maintenance of nasopharyngeal colonization by *S. aureus*. Notably, there was low maintenance of SNPs across passages, potentially indicating that selective pressures of each passage were distinct as the genetic variants exerted their phenotypes. Multiple contributing effects could also alter maintenance of sequence variants, including competitor microbiota, differences in host genetics of each host producing variations in the immune response or variation in the *S. aureus* community composition as the inoculated isolate pool replicates. The experimental design here, with sequential *in vivo* and *in vitro* steps could also contribute to the conservation of phenotype expression. These factors would account for both different mutations emerging, purifying selection and reversion of mutations. Alternatively, changes could indicate there was refinement in the repertoire of *S. aureus* genome variants during nasopharyngeal colonization. Observed selection outcomes would require that some mutations were dominant in their effects or could alternatively overcome reproduction limitation. There may also be multiple routes through selection of mutations to achieve the same broad aims. Determining the cause for the low SNP maintenance would require extensive assessment of the functional outcome of the mutations and mutation combinations.

The finding in this study that several polymorphisms led to amino acid changes identical with those identified in a previous *S. aureus* epidemiology study of epidemic community-associated methicillin-resistant *S. aureus* highlights there could be relevant functional outcomes of SNPs (Kennedy et al., 2008). Moreover, the overlap supports both the methodology used here to investigate relevant features of *S. aureus* nasopharyngeal colonization and further study of the shared polymorphisms.

Various genetic changes detected in our colonization study are relevant to metabolism and nucleic acid functions rather than virulence factors. Krismer et al. [37] determined that key metabolic pathways are essential during *S. aureus* nasal colonisation owing to the low nutrient levels. Mutations located in genes related to carbohydrate biosynthesis and transport (mannitol, sorbitol and fructose-6-P metabolism) were identified in our study that could impact upon the acquisition and use of alternative carbon sources during nasopharynx colonization. Further genetic variation was detected in genes for co-factors biotin and thiamine, linked to amino acid and carbohydrate metabolism. Chaffin et al. [38] reported increased *in vivo* transcription of co-factor genes associated with changes in the energy source and higher growth rate during lung infection. Young et al. [15] investigated *S. aureus* genetic changes from carriage to disease and found the majority of mutations were associated with regulation of persistence components instead of virulence factors. Mutations found in stress resistance genes in this study (*hemN, ahpF* and *clpL*) which are possibly linked to altered expression for *S. aureus* components involved in protection from oxidative stress in the anterior nares [39]. The absence of genetic variance selection in our study within virulence factors could reflect a predominant adaptive selection of metabolic pathways although the clear interplay in *S. aureus* between metabolism and virulence factor expression must not be overlooked [40].

Our identification of batches of mice pre-colonized with *S. aureus* proved to be a hindrance and future studies require that this effect is mitigated in the experimental plan. High rates of colonization are reported in these rodents from commercial vendors [25] and this appears to be the problem in our study, as none of the experimental *S. aureus* USA300 LAC JE2 inoculum was recovered from mice naturally pre-colonized with *S. aureus* SAU_MOU.. Both strains belonged to ST8 with different *spa* types (USA300 and SAU_MOU are t008 and t024, respectively). ST8 is an important clonal group, which includes the American epidemic CA-MRSA USA300 clone; this genotype is characterised by the unique linkage between SCC*mec*IVa and arginine catabolic mobile element (ACME), which provides both drug resistance and pronounced virulence factor expression during colonization/infection [41]. While pre-colonization of mice complicated completion of the experimental plan in our study, the outcomes supports the view that mice are better models for both colonization and infection than often argued [42].

Although USA300 LAC JE2 and SA_MOU share identical ST, genetic differences between these strains might account for the inability to identify that USA300 LAC JE2 had invaded the resident SA_MOU community colonized nasopharyngeal tissue to establish colonization, at least as judged by the lack of its retrieval. Genetic diversity within ST8 is known to be relatively high [29,43-45] and intraspecies genetic variation is known to prevent colonization by incoming *S. aureus* [28]. Votintseva et al. [46] used *spa* typing to detect multiple-strain *S. aureus* colonization and found that cocolonization was only 3.4 to 5.8% in samples pooled from healthy individuals.

In conclusion, our study highlights *S. aureus* genomic variants are selected using a model of nasopharynx colonization that incorporates successive inoculation cycles. The sequence variants were mostly within metabolism and stress pathways suggesting the associated gene functions could contribute to *S. aureus* nasal carriage. The individual understanding of their relevance will serve as basis for future studies focused on genetic factors that contribute to *S. aureus* nasal colonization and persistence.

## Materials and Methods

### Bacterial strains and growth conditions

The clinically-derived *S. aureus* strain USA300 LAC JE2 [47] was chosen as the strain to study in the model. This strain was used in the murine model described below which generated 90 nasopharyngeal colonization clones. Cultures were grown for 18 h at 37°C with shaking unless stated otherwise. Brain Heart Infusion (BHI) broth or agar was the medium used unless otherwise stated.

### Long-term nasopharyngeal colonization murine model

A novel and uniquelong-term nasopharyngeal colonization by S. aureus was established. Mouse experiments were approved by the United Kingdom Home Office (Home Office Project License Number 40/3602) and the University of Liverpool Animal Welfare and Ethics Committee.

Groups of three mice (female BALB/c mice of 6-8-week-old, Charles River, UK) were assigned to a cage and the small group size was based on the subsequent genetic analysis. Mice were maintained with food and water *ad libitum*.

Animals were anaesthetized with a mixture of oxygen and isofluorane and 5 × 10^4^ cfu of USA300 LAC JE2 were horizontally instilled into the nares. Mice were monitored over 14 days post-infection and did not show any visible signs of disease. At five time points over a period of 14 days, mice were culled by cervical dislocation followed by aseptic removal of nasopharynx to determine CFU load. No CFUs were recovered from the lungs or blood over this period. Nasopharyngeal tissue was mixed with 3 ml of PBS and mechanically disrupted using an IKA T10 handled tissue homogenizer. Bacteria were enumerated from viable counts of serially diluted nasopharyngeal homogenates with growth and species confirmation on mannitol salt agar after incubation for up to 48 h at 37 °C.

### Experimental evolution using nasopharyngeal colonization model

*Staphylococcus aureus* USA300 LAC JE2 was passaged three times by sequential colonization of the nasopharynx, with three mice per passage. Animal handling and bacterial inoculum preparation was performed as described above. An inoculum of USA300 LAC JE2 culture was prepared for the first nasopharyngeal passage and instilled in each of three mice. After the 7 d incubation period, mice were euthanized, each nasopharynx was aseptically removed and homogenized and the resulting tissue homogenates grown on separate mannitol salt agar plates. Ten clones were randomly selected from each replicate and subcultured. Equal volumes (1 ml) of the ten cultures were mixed to generate a new inoculum followed by dilution in PBS to obtain a bacterial load of ∼ 5 × 10^4^ cfu. This inoculum was used for a second passage of murine nasopharyngeal colonization. A third and final serial passage was achieved by repeating this approach. Both control and resulting clones from each of the three nasopharyngeal passages were stored alongside nasal homogenates at -80 °C with 15% glycerol (v/v).

### Genome Sequencing and analysis

DNA was purified from ten isolates randomly picked after plating nasal homogenates for each mouse from the three nasopharyngeal passages; purification methods were as described elsewhere [19] with the addition of an RNase step. The 9 genomic DNA pools for sequencing were prepared by mixing equimolar concentrations of DNA from which 9 libraries were prepared with an insert size of 350 bp using a TruSeq PCR-free sample prep kit (Illumina), according to the manufacturer’s instructions. DNA libraries were sequenced as barcoded pools on the Illumina MiSeq using V2 reagents generating paired sequencing reads of 350 bases in length. Reads were quality-filtered and trimmed for Nextera adaptor sequences using Trimmomatic [48]. Reads were aligned to parental *S. aureus* USA300 LAC JE2 that was sequenced in this study as a control. DNA library preparation and samples sequencing were done by the CGR (Centre for Genomic Research), University of Liverpool. Variant analysis was performed as described in Coates-Brown et al. [19] with a threshold of 20% of reads.

*S. aureus* strain SA_MOU isolated from mouse nasopharynx in this study was assembled then visualized using Geneious software (version R11.1) where the resulting assembly was aligned by BLASTZ software (version 7.0.1) to the USA300 reference genome (USA300_FPR3757; NCBI Ref Seq: NC_007793.1) and two additional reference genomes representing dominant STs in Europe: ST22 (HO 5096 0412; NCBI Ref Seq: NC_017763.1) and ST80 (NCTC 8325; NCBI Ref Seq: NC_007795.1).

### Sequencing of *spa* gene

Mouse nasopharynx isolates were screened by PCR amplification of the staphylococcal protein A (*spa*) gene to confirm that the *S. aureus* found in the nasopharynx was the same *spa* type as the inoculum strain. The reactions were undertaken using Mastercycler pro S (Eppendorf) and each contained 0.5 μM PA 1095F forward PCR primer, 0.5 μM PA 1517R reverse PCR primer [49], 80 ng DNA, 25 μL of BioMix Red (Bioline) and DEPC-treated water up to a total reaction size of 50 μL mixed into 0.2 mL PCR tubes. The run cycle parameters were an initial 10 min at 95°C and then 30 cycles of 30 s at 95°C, 30 s at 60°C and 45 s at °C with a final 10 min at 72°C. PCR products were purified by using the Isolate II PCR and gel kit (Bioline), analyzed by agarose gel electrophoresis and then sequenced (GATC Biotech) with samples prepared according to the company requirements. The *spa* sequence was analyzed by using BLASTn [50] with default settings. Matched PCR fragment sizes and BLASTn output data for both control and passaged clones were used to indicate that they belong to the same strain.

## Acknowledgments

DNA sequencing was performed by the Centre for Genomic Research, Liverpool.

## Data Availability Statement

The datasets generated for this study can be found in the ENA under accession number PRJEB43354.

## Competing Interests

*The authors declare that the research was conducted in the absence of any commercial or financial relationships that could be construed as a potential conflict of interest*.

## Financial Disclosure Statement

BABS was funded by the CAPES Foundation, Ministry of Education of Brazil. JCM was funded by BBRSC research grants BB/1532161 and BB/L023040/1. EMW and AK were funded by the Joint Programming Initiative on Antimicrobial Resistance (JPI-AMR). The funders were not involved in the study design, collection of samples, analysis of data or the writing of this report or the discussion to submit this report for publication.

## Supporting information

**S1 Table. Non-synonymous intragenic SNPs after one nasopharyngeal colonisation passage.**

**S2 Table. Intergenic SNPs after one nasopharyngeal colonisation passage**.

**S3 Table. Non-synonymous intragenic SNPs after two nasopharyngeal colonisation passages.**

**S4 Table. Intergenic SNPs after two nasopharyngeal colonisation passage**.

**S5 Table. Non-synonymous intragenic SNPs after three nasopharyngeal colonisation passages.**

**S6 Table. Intergenic SNPs after three nasopharyngeal colonisation passages**.

## Notes

### Competing Interest Statement

The authors have declared no competing interest.

