## Supplemental Tables 1 to 6 for "Long term nasopharyngeal colonization by *Staphylococcus aureus* determinants of adaptation"

**Table S1. Non-synonymous intragenic SNPs after one nasopharyngeal colonisation passage.**

*S. aureus* SNPs observed in pooled genomic DNA of ten isolates after the first passage of murine nasopharyngeal colonisation and sampling. Mouse isolate pools in which the SNPs were detected are indicated by A1, B1 and C1. Frequency in pool data shows the percentage of reads from each pool that harbours the SNP.

| **Gene symbol** | **Function** | **Genome change** | **Codon change** | **Mouse**  **Pool** | **Frequency in pool** |
| --- | --- | --- | --- | --- | --- |
| SAUSA300_RS00145 | Putative transposase in SCC*mec* element | 35982 (T/C) | 190 aa (L/F) | A1  C1 | 19.20%  21.51% |
| SAUSA300_RS02185 | Hypothetical protein  (SSL11) | 460966 (G/A) | 110 aa (A/T) | A1 | 21.45% |
| *valS* | Valyl-tRNA synthetase | 1762705 (T/C) | 800 aa (K/R) | C1 | 23.88% |

**Table S2. Intergenic SNPs after one nasopharyngeal colonisation passage.**

*S. aureus* SNPs observed in pooled genomic DNA of ten isolates after the first passage of murine nasopharyngeal colonisation and sampling. Mouse isolate pools in which the SNPs were detected are indicated by B1 and C1. Frequency in pool data shows the percentage of reads from each pool that harbours the SNP.

| **Genome change** | **Distance from coding region** | **Gene Function** | **Mouse**  **Pool** | **Frequency in pool** |
| --- | --- | --- | --- | --- |
| 1121360 (T/A) | -27 bp / SAUSA300_RS05520 (+)  -100 bp / SAUSA300_RS05515 (-) | Hypothetical protein  Hypothetical protein | A1  C1 | 19.20%  21.51% |

**Table S3. Non-synonymous intragenic SNPs after two nasopharyngeal colonisation passages.**

*S. aureus* SNPs observed in pooled genomic DNA of ten isolates after the second passage of murine nasopharyngeal colonisation and sampling. Mouse isolate pools in which the SNPs were detected are indicated by A2, B2 and C2, reflecting three independent mice. Frequency in pool data shows the percentage of reads from each pool that harbours the SNP.

| **Gene symbol** | **Function** | **Genome change** | **Codon change** | **Mouse**  **Pool** | **Frequency in pool** |
| --- | --- | --- | --- | --- | --- |
| *recF* | DNA recombination  and repair | 5010 (C/G) | 358 aa (P/A) | A2  B2  C2 | 66.08%  99.50%  89.47% |
| *gyrB* | DNA Topoisomerase II  subunit B | 5608 (T/C) | 183 aa (I/T) | A2  B2  C2 | 66.62%  99.87%  91.74% |
| *gyrA* | DNA Topoisomerase II  subunit A | 7282 (T/C) | 84 aa (L/S) | A2  B2  C2 | 67.60%  100%  92.43% |
| SAUSA300_RS00145 | Putative transposase in SCC*mec* element | 36124 (G/A) | 194 aa (A/V) | A2  B2  C2 | 35.68%  53.86%  51.40% |
| SAUSA300_RS00265 | Hypothetical  protein | 61025 (G/A) | 55 aa (L/F) | A2  B2  C2 | 63.37%  99.34%  93.45% |
| *cap5I* | Capsular PS  CP5 synthesis | 182746 (T/G) | 253 aa (L/V) | A2  B2  C2 | 66.755  98.75%  92.12% |
| SAUSA300_RS01070 | Hypothetical ABC transporter protein | 240358 (T/A) | 358 aa (N/K) | A2  B2  C2 | 66.15%  100%  92.37% |
| SAUSA300_RS01200 | Hydroxyacyl Coenzyme A dehydrogenase | 270510 (T/C) | 601 aa (T/A) | A2  B2  C2 | 62.76%  97.07%  90.58% |
| *gutB* | Sorbitol  dehydrogenase | 292738 (A/C) | 341 aa (T/P) | A2  B2  C2 | 64.99%  99.64%  91.64% |
| SAUSA300_RS01310 | Cell wall and capsule  metabolism | 294913 (G/A) | 119 aa (G/D) | A2  B2  C2 | 64.14%  99.77%  92.05% |
| SAUSA300_RS01715 | Oye family flavin  oxidoreductase | 376057 (C/T) | 258 aa (D/N) | A2  B2  C2 | 65.77%  99.86%  92.24% |
| SAUSA300_RS01715 | Oye family flavin  oxidoreductase | 376231 (T/G) | 200 aa (S/R) | A2  B2  C2 | 63.97%  99.73%  92.89% |
| *ahpF* | Alkyl hydroperoxide  reductase | 429073 (C/T) | 108 aa (G/D) | A2  B2  C2 | 64.37%  100%  91.73% |
| SAUSA300_RS02220 | Hypothetical lipoprotein  (Lpl4) | 467549 (A/T) | 209 aa (E/D) | A2  B2  C2 | 68.29%  99.26%  89.27% |
| SAUSA300_RS03590 | Undecaprenyl diphosphatase | 742477 (C/T) | 69 aa (R/H) | A2  B2  C2 | 69.23%  99.58%  90.45% |
| SAUSA300_RS04050 | Sporulation Regulator  WhiA | 837108 (G/T) | 17 aa (E/D) | A2  B2  C2 | 67.52%  100%  93.17% |
| *ampA* | Cytosol  aminopeptidase | 923896 (G/T) | 207 aa (K/A) | A2  B2  C2 | 64.95%  99.72%  91.94% |
| SAUSA300_RS04890 | Pseudouridine  synthase | 996224 (C/T) | 150 aa (S/F) | A2  B2  C2 | 65.68%  100%  91.67% |
| SAUSA300_RS05275 | Hypothetical protein | 1074815 (G/A) | 71 aa (D/N) | A2  B2  C2 | 64.97%  99.75%  92.06% |
| *sun* | 16S rRNA  methyltransferase | 1215913 (T/C) | 306 aa (Y/H) | A2  B2  C2 | 64.19%  99.87%  90.09% |
| *ftsK* | DNA translocase | 1286986 (A/G) | 180 aa (K/R) | A2  B2  C2 | 65.14%  100%  91.76% |
| SAUSA300_RS06375 | RNase Y | 1298198 (T/G) | 28 aa (D/E) | A2  B2  C2 | 68.71%  99.85%  93.01% |
| *sbcC* | DNA repair: exonuclease | 1363207 (A/T) | 761 aa (I/F) | A2  B2  C2 | 66.24%  99.84%  91.75% |
| *parC* | Toposiomerase IV subunit A | 1374405 (A/C) | 80 aa (T/S) | A2  B2  C2 | 63.75%  99.85%  90.89% |
| SAUSA300_RS07265 | Hypothetical protein | 1496455 (T/A) | 663 aa (I/L) | A2  B2  C2 | 65.75%  99.86%  93.06% |
| SAUSA300_RS07925 | SDR family  Oxidoreductase | 1604107 (T/C) | 196 aa (E/G) | A2  B2  C2 | 61.10%  100%  92.77% |
| SAUSA300_RS08410 | Copro-porphyrinogen  III oxidase | 1693739 (T/C) | 145 aa (Y/A) | A2  B2  C2 | 67.43%  100%  91.83% |
| SAUSA300_RS09205 | Hypothetical protein | 1857089 (C/T) | 179 aa (V/I) | B2  C2 | 23.98%  20.28% |
| SAUSA300_RS09205 | Hypothetical protein | 1857109 (A/G) | 172 aa (I/T) | B2  C2 | 29.15%  24.40% |
| SAUSA300_RS09205 | Hypothetical protein | 1857182 (G/A) | 148 aa (P/S) | A2  B2  C2 | 31.10%  44.48%  40.93% |
| SAUSA300_RS09205 | Hypothetical protein | 1857188 (C/T) | 146 aa (E/K) | A2  B2  C2 | 31.83%  47.30%  42.20% |
| SAUSA300_RS09205 | Hypothetical protein | 1857202 (G/A) | 141 aa (A/V) | A2  B2  C2 | 31.97%  42.25%  41.06% |
| SAUSA300_RS09205 | Hypothetical protein | 1857215 (T/C) | 137 aa (T/A) | A2  B2  C2 | 35.55%  51.15%  46.06% |
| SAUSA300_RS09205 | Hypothetical protein | 1857232 (T/C) | 131 aa (K/R) | A2  B2  C2 | 39.91%  56.41%  52.01% |
| SAUSA300_RS09220 | FtsK/SpoIIIE  Family protein | 1859999 (C/T) | 1142 aa(G/D) | A2  B2  C2 | 65.07%  99.86%  90.30% |
| SAUSA300_RS09290 | Hypothetical  flavoprotein | 1876812 (A/T) | 97 aa (D/V) | A2  B2  C2 | 65.61%  100%  91.97% |
| SAUSA300_RS09425 | Transaldolase | 1908169 (T/C) | 6 aa (I/V) | A2  B2  C2 | 67.73%  99.87%  90.37% |
| SAUSA300_RS09885 | Epoxyqueuosine  reductase | 1988983 (C/T) | 174 aa (V/I) | A2  B2  C2 | 66.57%  99.73%  90.37% |
| SAUSA300_RS10590 | Phage tail tape measure protein | 2100157 (A/C) | 384 aa (I/S) | A2  B2  C2 | 674.75%  100%  92.87% |
| SAUSA300_RS10650 | Phage terminase protein | 2109096 (G/T) | 84 aa (A/D) | A2  B2  C2 | 74.13%  99.73%  91.26% |
| SAUSA300_RS11285 | Thiaminase II | 2214645 (C/T) | 224 aa (G/R) | A2  B2  C2 | 63.53%  100%  91.84% |
| *mtlA* | PTS system Mannitol transporter | 2276564 (A/G) | 56 aa (E/G) | A2  B2  C2 | 65.21%  99.75%  89.40% |
| *narK* | Nitrite extrusion  protein | 2508817 (G/A) | 241 aa (A/V) | A2  B2  C2 | 66.90%  100%  90.41% |
| SAUSA300_RS13805 | Clp protease ATP-binding subunit | 2685939 (T/G) | 249 aa (S/A) | A2  B2  C2 | 66.21%  100%  90.81% |
| SAUSA300_RS13825 | Transport protein | 2690087 (T/G) | 795 aa (K/G) | A2  B2  C2 | 63.58%  99.66%  94.32% |
| *hisG* | ATP phosphoribosyl-  transferase | 2839280 (C/A) | 140 aa (G/C) | A2  B2 | 22.62%  31.12% |

**Table S4. Intergenic SNPs after two nasopharyngeal colonisation passage.**

*S. aureus* genome sequence variants (SNPs) in non-coding regions observed for isolates after the second passage performed in the experimental nasopharyngeal evolution by. Mouse column relates to the pool of isolates (A2, B2 and C2) which the SNPs were detected. Frequency shows the percentage of reads from the pools harbouring the SNP.

| **Genome change** | **Distance from coding region** | **Gene Function** | **Mouse**  **Pool** | **Frequency in pool** |
| --- | --- | --- | --- | --- |
| 35817* (A/T) | sRNA sequence (Teg5as) |  | A2  B2  C2 | 45.98%  56.945  57.29% |
| 74496 (T/G) | -1069 bp / SAUSA300_RS00340 (+)  -148 bp (arcA) (-) | Universal stress protein  Arginine deiminase | A2  B2  C2 | 62.18%  99.77%  92.38% |
| 260357 (C/T) | -222 bp / *pflB* (+)  -366 bp / *hptA* (-) | Acetyltransferase  Iron-binding protein | A2  B2  C2 | 67.00%  100%  91.01% |
| 349622 (T/C) | -517 bp / SAUSA300_RS01590 (+) | Hypothetical Protein | A2  B2  C2 | 62.02%  99.79%  91.61% |
| 585363 (G/A) | -191 bp / *rpoB* (+) | DNA-directed RNA polymerase | A2  B2  C2 | 67.96%  100%  91.28% |
| 940710 (A/G) | -9 bp / SAUSA300_RS15240 (+)  -18 bp / SAUSA300_RS15245 (-) | Pseudogene  Hypothetical protein | A2  B2  C2 | 54.30%  99.73%  85.03% |
| 940718 (A/G) | -17 bp / SAUSA300_RS15240 (+)  -10 bp / SAUSA300_RS15245 (-) | Pseudogene  Hypothetical protein | A2  B2  C2 | 56.38%  100%  85.80% |
| 1161534 (C/T) | -44 bp / *argF* (+)  -394 bp / SAUSA300_RS05750 (-) | Amino acid metabolism  Superantigen-like protein | A2  B2  C2 | 68.70%  100%  93.17% |
| 1533312 (A/T) | -237 / SAUSA300_RS07455 (+)  -165 bp / *rpsA* (-) | Hypothetical protein  30S ribosomal protein S1 | A2  B2  C2 | 66.52%  99.86%  91.07% |
| 1929046 (T/C) | -289 bp / SAUSA300_RS09565 (+)  -132 bp / SAUSA300_RS09560 (-) | Hypothetical protein  Hypothetical transposase | A2  B2  C2 | 45.10%  54.13%  57.68% |
| 1929047 (G/A) | -288 bp / SAUSA300_RS09565 (+)  -133 bp / SAUSA300_RS09560 (-) | Hypothetical protein  Hypothetical transposase | A2  B2  C2 | 45.04%  54.30%  57.55% |
| 1929049 (C/A) | -286 bp / SAUSA300_RS09565 (+)  -135 bp / SAUSA300_RS09560 (-) | Hypothetical protein  Hypothetical transposase | A2  B2  C2 | 45.08%  54.10%  57.32% |
| 1929054 (A/G) | -280 bp / SAUSA300_RS09565 (+)  -141 bp / SAUSA300_RS09560 (-) | Hypothetical protein  Hypothetical transposase | A2  B2  C2 | 45.72%  54.78%  58.31% |
| 1929062 (G/A) | -272 bp / SAUSA300_RS09565 (+)  -149 bp / SAUSA300_RS09560 (-) | Hypothetical protein  Hypothetical transposase | A2  B2  C2 | 47.73%  56.78%  59.75% |
| 1929098* (C/A) | sRNA sequence (SprA1) |  | A2  B2  C2 | 46.37%  55.89%  58.84% |
| 1929167* (C/T) | sRNA sequence (SprA1) |  | A2  B2  C2 | 45.53%  55.43%  57.57% |
| 1929168* (A/G) | sRNA sequence (SprA1) |  | A2  B2  C2 | 45.52%  55.48%  57.50% |
| 1929175* (T/A) | sRNA sequence (SprA1) |  | A2  B2  C2 | 45.64%  56.03%  57.39% |
| 1929178* (T/A) | sRNA sequence (SprA1) |  | A2  B2  C2 | 45.745  56.01%  57.18% |
| 1929180* (T/A) | sRNA sequence (SprA1) |  | A2  B2  C2 | 46.14%  56.09%  57.43% |
| 1929191* (A/T) | sRNA sequence (SprA1) |  | A2  B2  C2 | 44.82%  55.04%  56.90% |
| 1929196* (G/A) | sRNA sequence (SprA1) |  | A2  B2  C2 | 45.36%  54.93%  57.04% |
| 1957938 (C/T) | -100 bp / SAUSA300_RS09690 (+)  -699 bp / *lukE* (-) | Hypothetical protein  Leukotoxin | A2  B2  C2 | 65.46%  99.88%  92.30% |
| 1959504 (T/A) | -313 bp / SAUSA300_RS15420 (+) | Pseudogene | A2  B2  C2 | 66.46%  100%  90.40% |
| 2322700 (T/C) | -290 bp / *opuD2* (-) | ABC transporter permease | A2  B2  C2 | 67.73%  100%  91.06% |

*SNPs located in regulatory sRNA sequence

**Table S5. Non-synonymous intragenic SNPs after three nasopharyngeal colonisation passages.**

*S. aureus* genome sequence variants (SNPs) in non-coding regions observed for isolates after the third passage in an experimental nasopharyngeal model. Mouse column relates to the pool of isolates (A3, B3 and C3) which the SNPs were detected. Frequency shows the percentage of reads from the pools harbouring the SNP.

| **Gene symbol** | **Function** | **Genome change** | **Codon change** | **Mouse**  **Pool** | **Frequency in pool** |
| --- | --- | --- | --- | --- | --- |
| *recF* | DNA replication and repair | 5010 (C/G) | 358 aa (P/A) | A3  B3  C3 | 100%  100%  100% |
| SAUSA300_RS00265 | Hypothetical protein | 61025 (G/A) | 55 aa (L/F) | A3  B3  C3 | 99.97%  100%  99.98% |
| SAUSA300_RS01070 | RGD-containing  lipoprotein | 240358 (T/A) | 358 aa (N/K) | A3  B3  C3 | 100%  100%  100% |
| SAUSA300_RS01200 | Hydroxyacyl Coenzyme A dehydrogenase | 270510 (T/C) | 601 aa (T/A) | A3  B3  C3 | 99.98%  100%  100% |
| *gutB* | Sorbitol dehydrogenase | 292738 (A/C) | 341 aa (T/P) | A3  B3  C3 | 99.98%  100%  100% |
| SAUSA300_RS01715 | Oye family flavin  oxireductase | 376057 (C/T) | 258 aa (D/N) | A3  B3  C3 | 99.98%  99.93%  99.97% |
| SAUSA300_RS01715 | Oye family flavin oxireductase | 376231 (T/G) | 200 aa (S/R) | A3  B3  C3 | 99.97%  100%  100% |
| SAUSA300_RS02185 | Staphylococcal superantigen-like 11 | 460966 (G/A) | 110 aa (A/T) | A3  B3  C3 | 100%  92.30%  95.83% |
| *ftsK* | DNA translocase | 1286986 (A/G) | 180 aa (K/R) | A3  B3  C3 | 100%  100%  100% |
| *dnaJ* | Chaperone protein | 1688528 (C/T) | 250 aa (A/T) | A3  B3  C3 | 99.97%  100%  99.95% |
| SAUSA300_RS09205 | Hypothetical protein | 1857089 (C/T) | 179 aa (V/I) | B3 | 13.10% |
| SAUSA300_RS09205 | Hypothetical protein | 1857109 (A/G) | 172 aa (I/T) | A3  B3  C3 | 17.90%  24.42%  24.67% |
| SAUSA300_RS09205 | Hypothetical protein | 1857182 (G/A) | 148 aa (P/S) | A3  B3  C3 | 38.13%  41.51%  41.87% |
| SAUSA300_RS09205 | Hypothetical protein | 1857188 (C/T) | 146 aa (E/K) | A3  B3  C3 | 41.28%  43.10%  43.85% |
| SAUSA300_RS09205 | Hypothetical protein | 1857202 (G/A) | 141 aa (A/V) | A3  B3  C3 | 40.21%  40.75%  40.88% |
| SAUSA300_RS09205 | Hypothetical protein | 1857215 (T/C) | 137 aa (T/A) | A3  B3  C3 | 46.58%  45.50%  45.44% |
| SAUSA300_RS09205 | Hypothetical protein | 1857232 (T/C) | 131 aa (K/R) | A3  B3  C3 | 50.28%  49.51%  50% |
| SAUSA300_RS15385 | Hypothetical protein | 1872844 T/C) | 28 aa (Y/C) | A3  B3  C3 | 99.97%  99.95%  99.94% |
| SAUSA300_RS10080 | Hypothetical protein | 2007223 (G/T) | 48 aa (R/L) | A3  B3  C3 | 100%  99.97%  99.96% |
| *bioA* | Aminotransferase | 2549677 (C/T) | 189 aa (R/H) | A3  B3  C3 | 99.96%  99.93%  99.98% |

**S6 Table. Intergenic SNPs after three nasopharyngeal colonisation passages.**

*S. aureus* genome sequence variants (SNPs) in non-coding regions observed for isolates after the third passage in an experimental nasopharyngeal model. Mouse pool column relates to the pool of isolates (A3, B3 and C3) which the SNPs were detected. Frequency shows the percentage of reads from the pools harbouring the SNP.

| **Genome change** | **Distance from coding region** | **Gene Function** | **Mouse**  **Pool** | **Frequency in pool** |
| --- | --- | --- | --- | --- |
| 164912 (G/A) | -97 bp / SAUSA300_RS00765 (+)  -132 bp / *phnD* (-) | Hypothetical protein  Phosphonate transporter | A3  B3  C3 | 45.98%  56.94%  57.29% |
| 1533312 (A/T) | -237 / SAUSA300_RS07455 (+)  -165 bp / *rpsA* (-) | Hypothetical protein  30S ribosomal protein S1 | A3  B3  C3 | 99.98%  99.94%  99.94% |
| 1957938 (C/T) | -100 bp / SAUSA300_RS09690 (+)  -699 bp / *lukE* (-) | Hypothetical protein  Leukotoxin | A3  B3  C3 | 99.97%  100%  99.98% |
| 2164596* (A/G) | sRNA sequence (Teg16) |  | A3  B3  C3 | 62.18%  99.77%  92.38% |
| 2322700 (A/G) | -290 bp / *opuD2* (-) | ABC transporter permease | A3  B3  C3 | 100%  100%  99.98% |
| 2487639 (A/G) | -189 bp / SAUSA300_RS12785 (+)  -128 bp / *lctP2* (-) | Phage transferase  Lactate permease | A3  B3  C3 | 67.00%  100%  91.01% |

*SNPs located in regulatory sRNA sequence
